# Using the Phenoscape Knowledgebase to relate genetic perturbations to phenotypic evolution

**DOI:** 10.1101/018853

**Authors:** Prashanti Manda, James P. Balhoff, Hilmar Lapp, Paula Mabee, Todd J. Vision

## Abstract

The abundance of phenotypic diversity among species can enrich our knowledge of development and genetics beyond the limits of variation that can be observed in model organisms. The Phenoscape Knowledgebase (KB) is designed to enable exploration and discovery of phenotypic variation among species. Because phenotypes in the KB are annotated using standard ontologies, evolutionary phenotypes can be compared with phenotypes from genetic perturbations in model organisms. To illustrate the power of this approach, we review the use of the KB to find taxa showing evolutionary variation similar to that of a query gene. Matches are made between the full set of phenotypes described for a gene and an evolutionary profile, the latter of which is defined as the set of phenotypes that are variable among the daughters of any node on the taxonomic tree. Phenoscape’s semantic similarity interface allows the user to assess the statistical significance of each match and flags matches that may only result from differences in annotation coverage between genetic and evolutionary studies. Tools such as this will help meet the challenge of relating the growing volume of genetic knowledge in model organisms to the diversity of phenotypes in nature. The Phenoscape KB is available at http://kb.phenoscape.org.

## Introduction

Millions of years of evolution have led to a vast range of phenotypic diversity among and within species, and a correspondingly large literature on comparative anatomy and morphological systematics. While the literature on a particular taxon can be mastered by an individual expert, it is very challenging to discover what is known about variation in a particular phenotype across distantly related taxa. This limits our ability to apply discoveries from developmental genetics to natural variation, and to take advantage of naturally occurring phenotypic variation for enriching our understanding how organisms are built.

In recent years, the Phenoscape project (phenoscape.org) has been working to enable discovery of what is known about natural variation in phenotypes from the literature. Here, we explain how researchers can use Phenoscape resources to discover natural phenotypic variation that bears similarity to that seen when perturbing a particular gene in a model organism.

## Phenoscape

Phenotypes in the evolutionary literature are typically described in natural language (Dahdul et al., 2010). While wonderfully expressive, such descriptions are not easily integrated across different studies and taxa using computational tools. A shared, consistent, and machine-readable representation of phenotypes is key for enabling computer-aided exploration, knowledge discovery, and hypothesis generation from heterogeneous phenotypes (Deans et al., 2015).

Towards this goal, the Phenoscape project has spearheaded the development of multispecies anatomy ontologies (Dahdul et al. 2010b; 2012), has contributed to community ontologies (Haendel et al., 2014; Gkoutos et al., 2005; Midford et al. 2013), developed data annotation software (Balhoff et al., 2010; Balhoff et al., 2014), developed methodology for annotating evolutionary phenotypes from the literature (Dahdul et al. 2010a), and built the Phenoscape Knowledgebase (KB), an ontology-driven database that combines existing phenotype annotations from model organism databases with new phenotype annotations from the evolutionary literature. The contents of the KB at time of writing KB (2015-Apr-29) are summarized in **Table 1**, and described in more detail in the following sections.

**Table 1.** Summary of phenotype and expression data in the Phenoscape Knowledgebase (as of 2015-Apr-29).

**A. Evolutionary data**
|  |  |
| --- | --- |
| Annotated anatomical character states | 19,547 |
| Total number of annotated taxa (vertebrates) | 4,895 |
| Terminal taxa with at least one phenotype | 3,935 |
| Non-terminal taxa with at least one phenotype | 960 |
| Evolutionary phenotype profiles | 636 |

**B. Model organism data**
|  | Zebrafish | Mouse | Xenopus | Human |
| --- | --- | --- | --- | --- |
| Genes with at least one phenotype | 5,109 | 7,758 | 12 | 3,050 |
| Phenotype annotations | 68,433 | 171,876 | 236 | 80,044 |
| Genes with any expression data* | 11,832 | 10,599 | 15,030 | 0 |
| Gene expression annotations | 160,255 | 800,824 | 328,324 | 0 |

## Model organism phenotypes

Gene phenotypes are phenotypic observations made using knockouts, induced mutants, overexpression constructs and other single-gene perturbations, and are described relative to a wild-type phenotype. A gene phenotype profile consists of the union of all the single-gene phenotypes observed across different alleles, backgrounds, and types of perturbations (knockouts, over-expression constructs, etc.) for each model organism. Thus, there are distinct gene profiles for orthologous genes in different taxa. Gene phenotype profiles in the Phenoscape KB are from mouse (MGI, Eppig et al., 2015), *Xenopus* (Xenbase, Karpinka et al., 2015), zebrafish (ZFIN, Bradford et al., 2011), and human (Human Phenotype Ontology project, Kohler et al., 2013). Though not used in the semantic similarity search described here, the KB also includes information on the gross anatomical localization of expression for genes, imported from Xenbase and ZFIN in the case of *Xenopus* and zebrafish, respectively, and from MGI GXD in the case of mouse (Smith et al., 2014).

## Evolutionary phenotypes

The Phenoscape KB currently contains, and is the original source for, phenotype annotations for more than 4,800 species and higher taxa, including over 19,000 character states sourced from 149 published phylogenetic studies. Each character state has been translated into one or more ontology-based Entity–Quality (EQ) annotations (Mungall et al., 2010) using the Phenex software (Balhoff et al., 2010) following the curation process in Dahdul et al. (2010). The EQ annotations are frequently less detailed than the original free-text description. Entities are taken from the Uberon anatomy ontology (Mungall et al., 2012; Haendel et al., 2014), qualities from the Phenotype and Trait Ontology (PATO, Gkoutos et al., 2005), and taxa from the Vertebrate Taxonomy Ontology (VTO, Midford et al., 2013).

Character states, and corresponding annotations, are initially annotated to individual taxa, and about 80% of the time those are terminal taxa, (*i.e.* species) (**Table 1**). Because the character states are taken from phylogenetic studies, they are phylogenetically informative, meaning that they are shared among closely related species but variable among the full set of species included in the study, due to character evolution in some ancestor. An inference step is necessary to infer where in the evolutionary history of the lineage the phenotypic change has occurred, based on the pattern of variation among descendant taxa. There are sophisticated methods for making such inferences that account for a variety of uncertainties and for different kinds of characters (Felsenstein 2004). The Phenoscape KB uses a simple approach for discrete changes based on the parsimony algorithm of Fitch (1971). First, the set of character states for a taxon is defined to include the character states annotated to all descendant taxa. Then, characters with sets of states that differ among the daughter lineages of a particular taxon are noted as being variable for that taxon. The evolutionary phenotype profile of a taxon consists of the set of all ontology annotations for characters that vary in state among the immediate descendants of that taxon. These evolutionary profiles are precomputed within the KB.

To take an example, the genus *Leporellus* (a so-called ‘headstander fish’), has four daughter species (**Figure 1**). Two of these, *L. pictus* and *L. vittatus,* are associated with 158 and 159 character states respectively, while the other two have none. Despite the large number of characters in *L. pictus* and *L. vittatus*, only two differ in state between them: the “number of cusps on second and third teeth of premaxilla” and the “form of cusping of teeth on fifth upper pharyngeal tooth-plate” (**Table 2**). Thus, the evolutionary phenotype profile of *Leporellus* consists of the EQ annotation(s) that correspond to these two characters.

**Table 2.**
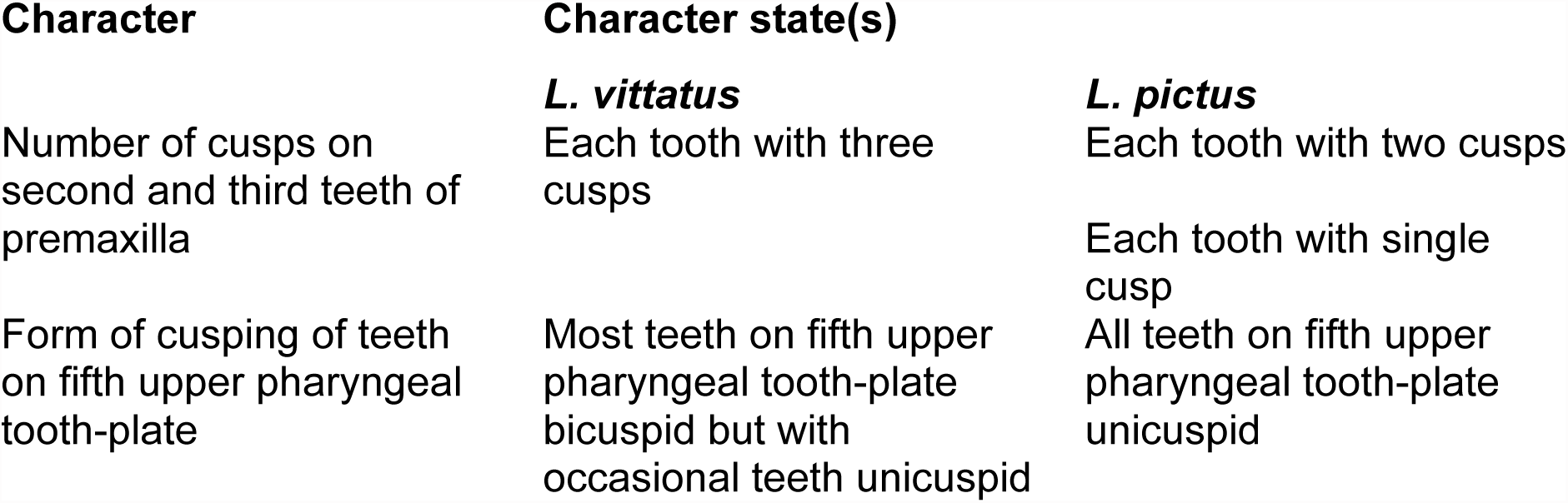
Characters with states that vary between two daughter taxa of the fish genus *Leporellus*, *L. vittatus* and *L. pictus.*

**Figure 1.**
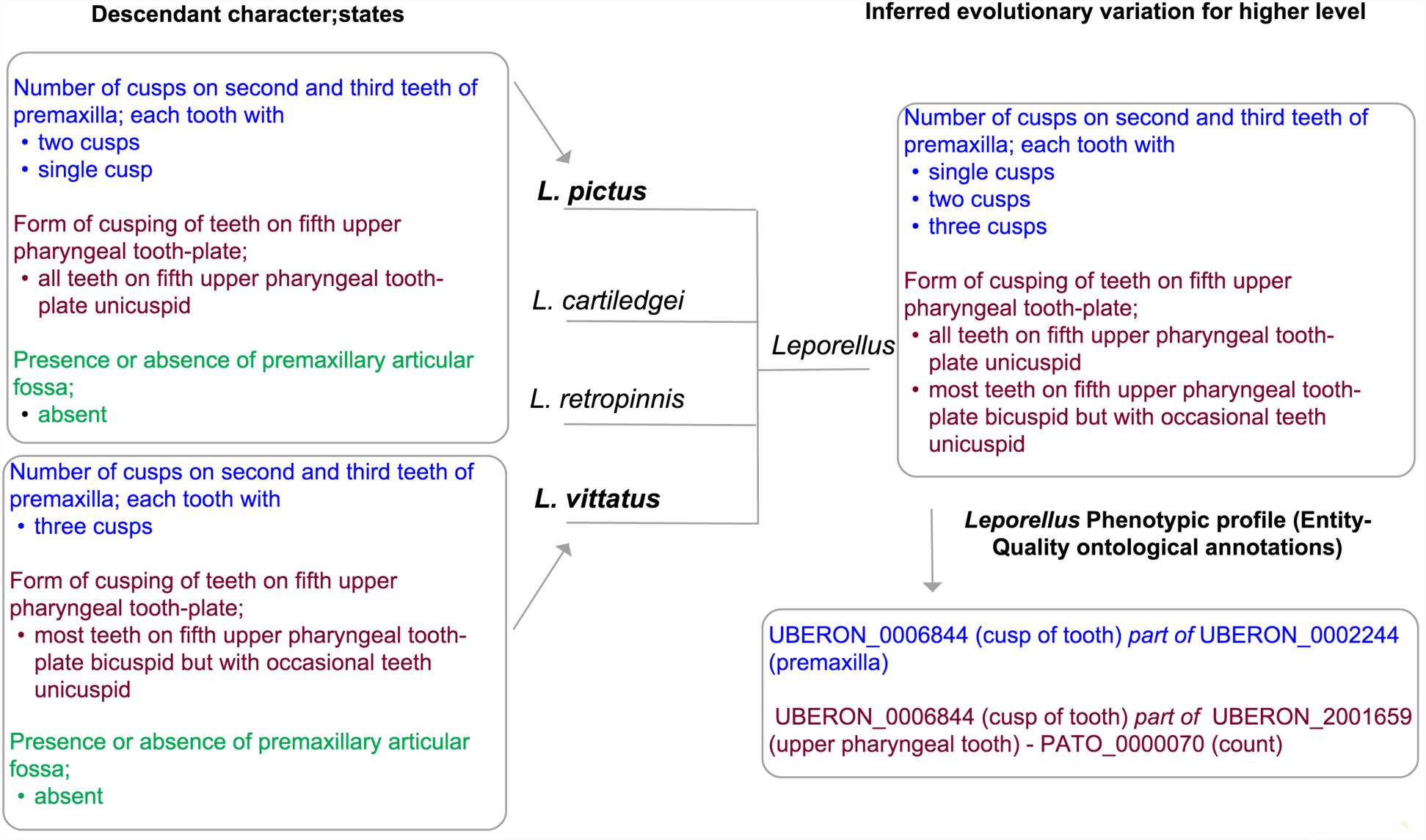
Inference of an evolutionary profile. Of the four species immediately descended from genus *Leporellus*, two (in bold) have phenotype annotations, a subset of which is shown in the left-hand boxes. Each annotation contains a character and one or more states separated by a semicolon. The two characters that vary in their states (blue and maroon) contribute to the evolutionary profile of the parent taxon while the one that is invariant (green) does not. At lower right, the evolutionary profile of *Leporellus* is translated into a set of ontological annotations that can be used for semantic similarity matching.

It is important to recognize the limitations to the inference of evolutionary phenotype profiles. Ideally, one would like to group phenotypic changes that are either pleiotropic, or at least occurred at a single point of evolutionary time. However, evolutionary phenotype profiles may group phenotypes that are genetically and evolutionarily independent. This is in part due to the branching patterns in the taxonomy, which does not resolve the relative order of branching within a taxon, and may be inaccurate relative to the true phylogeny. Even if the true, fully resolved, phylogeny were known, the parsimony algorithm may place the evolutionary change at an incorrect position (Cunningham et al. 1999). Finally, even if the ancestral reconstruction is correct, two phenotypes that show variation among the descendants of a given taxon may be the result of independent genetic changes.

It is also important to recognize that, due to the manual effort involved in curation (Dahdul et al., 2015), the KB necessarily includes a very incomplete sample of evolutionary phenotypes in the vertebrates, with coverage varying across taxonomic groups and anatomical systems. Across the skeleton, coverage is strongest in ostariophysan fishes, the group that includes zebrafish. Coverage is also good for fin, limb and girdle phenotypes from basal sarcopterygian fishes to early amphibians, because of the interest in phenotypes potentially related to the fin-to-limb transition from fishes to tetrapods (Mabee et al., 2012). Nonetheless, the absence of phenotype variation needs to be interpreted with care, and interpretation depends on the taxon and character.

## Semantic similarity

An important application of the KB is to enable discovery of evolutionary phenotype profiles that are more similar than one would expect by chance to a given gene phenotype profile. This application is similar in many ways to the popular BLAST tool, used to search large databases for similarity in nucleotide or protein sequence (Altschul et al., 1997). In our case however, the common ontologies used for gene phenotypes and evolutionary phenotypes allow one to measure *semantic similarity* (Pesquita et al., 2009). Semantic similarity allows for imperfect matches both at the level of individual EQ statements and for profiles as a whole.

Model organism gene phenotypes and evolutionary phenotypes used in the similarity search are described using a variety of ontologies and formats requiring a comprehensive ontology that connects these disparate ontologies. Moreover, the EQ annotation format post-composes concepts (Mungall et al., 2010) from multiple ontologies, requiring additional subsumer classes to enable reasoning in the absence of a pre-composed EQ ontology. A comprehensive ontology was compiled for the similarity search by incorporating individual ontologies used by model organism communities along with appropriate interconnecting “bridges”. In addition, approximately 288,000 ontology classes representing all possible combinations of anatomical entities (from Uberon) and quality attribute classes (such as ‘shape’, ‘count’ etc. from PATO) were added to the comprehensive ontology for reasoning over EQ annotations.

The semantic similarity measure used by the Phenoscape KB is based on the Information Content (IC, Resnik, 1999) of the most informative common ancestor (MICA) in the ontology between two phenotypes. The IC score is inversely proportional to the frequency of annotations to that MICA among all the evolutionary profiles in the KB, such that matches are scored more strongly when their commonality is both rare, and therefore assumed to be specific. **Figure 2** shows an example of a MICA for a single phenotype from the gene phenotype profile for *eda* in zebrafish and a single phenotype from the evolutionary phenotype profile for the genus *Leporellus*. The fewer evolutionary profiles that include a phenotype subsumed by ‘*part of* pharyngeal arch 7 skeleton - count’ and its descendants, the higher the IC score would be for this match, i.e., the MICA. To compute the overall score between two profiles, the KB uses the median score among all pairwise matches between the constituent EQ annotations of the gene phenotype profile and those of the taxon phenotype profile. All Match IC scores and the overall similarity scores are reported in normalized form, divided by the maximum possible score, so that they range from zero to one.

**Figure 2.**
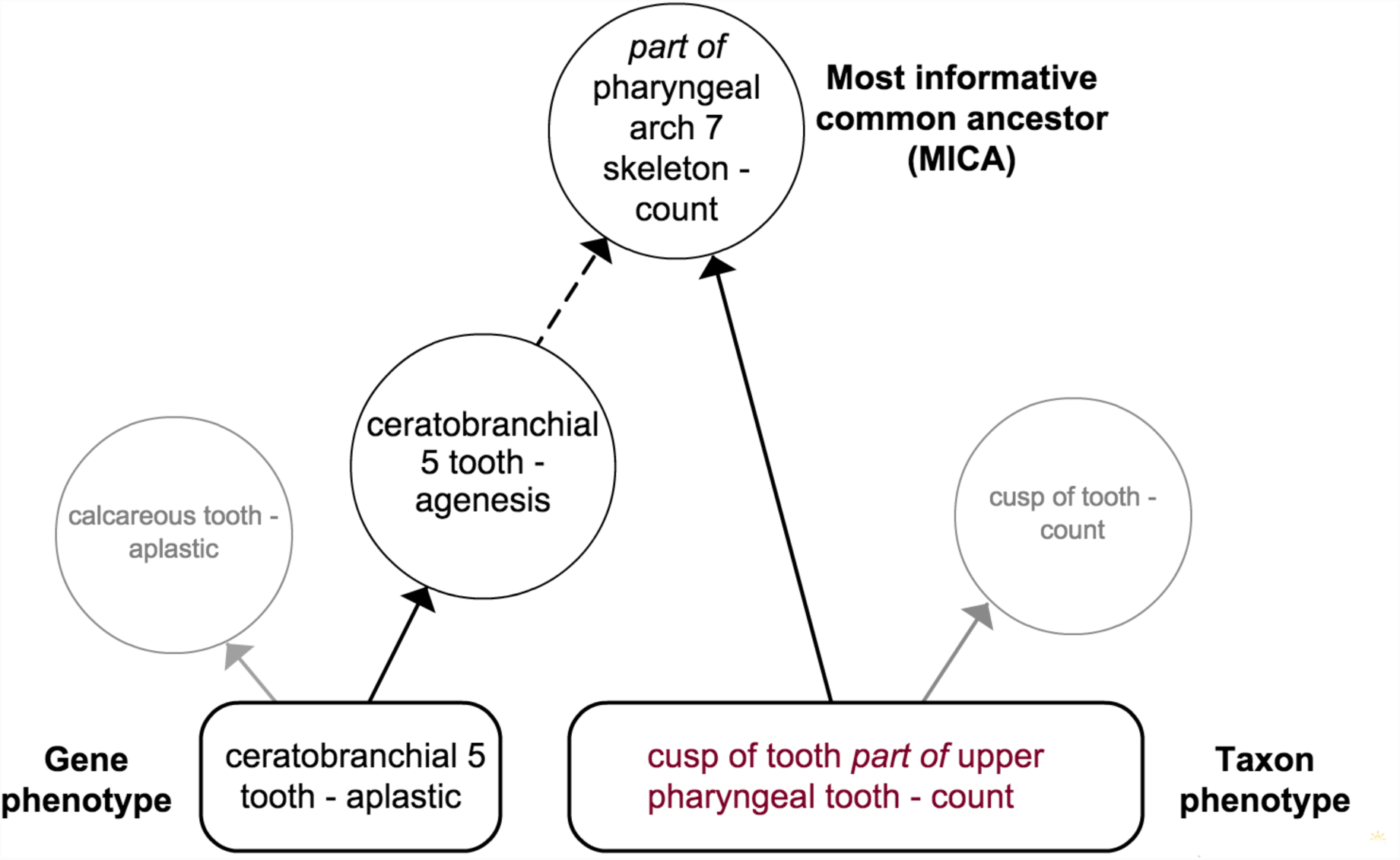
A match between ontologically annotated gene and taxon (evolutionary) phenotypes. One of the phenotypes from the profile of the *eda* gene in zebrafish matches one of the phenotypes from the evolutionary profile of *Leporellus* at the subsuming class ‘*part of* pharyngeal arch 7 skeleton – count’. This is the most informative common ancestor (MICA), though many other matches are possible (e.g., *part of* tooth – count). Solid arrows indicate direct subclass relationships whereas broken arrows indicate that one or more classes on the path have been hidden for clarity. The Information Content (IC) of the MICA is taken to be the similarity score for the phenotype pair.

To evaluate whether the score obtained is greater than would be expected by chance, the Phenoscape KB reports an Expect (*E*) value, which can be interpreted as the number of evolutionary profiles one would expect to see for a given gene profile with the same median IC or higher. It is based on a multiple linear regression of similarity scores against the two profile sizes. Studentized residuals from the regression are converted to *E*-values using Equations 3 and 4 from Pearson (1998) and adjusted for the size of the database being searched. *E*-values can range from 0 to the maximum number of taxa with evolutionary profiles in the KB, which is 636 in the current version, but one should consider there to be good evidence for a match only when the *E*-value is substantially less than one.

For some matches between individual phenotypes, the KB displays a warning when the MICA has a substantially higher IC among the evolutionary phenotype profiles than if the calculation had been done using the frequencies of annotations among the gene phenotype profiles. Phenotype pairs flagged in this way are expected to contribute high-scoring matches for many genes to this same evolutionary profile, and so the user may wish to discount them as evidence for a strong match between the two profiles. By default, a pair is flagged when the difference is at least 0.25.

### An example: *Wnt7a*

The Phenoscape KB enables users to query for evolutionary phenotype profiles that match any gene for which there is a genetic phenotypic profile available from mouse, human, zebrafish, or *Xenopus*. To illustrate this functionality, we will use the mouse gene *Wnt7a,* which has been hypothesized to be a candidate in the fin to limb evolutionary transition (Gibson-Brown et al., 1996; Hinchliffe et al., 2002; Yano et al., 2013; Abbasi 2013).

The user begins by entering the first few letters of a gene name in the auto-completing search bar (**Figure 3A**) and selecting the gene from the model organism of interest from the drop down list of the gene in different model organism species. When *Wnt7a* (mouse) is selected, the KB displays the Summary View listing the top-matching evolutionary phenotype profiles ranked by Expect Score (*E*-value) (**Figure 3B**). Here, the most similar match to *Wnt7a* is the suborder Microchiroptera (bats) with an *E*-value of 0.06. For each match in the Summary View, a Detail View is available (**Figure 3C**) which shows the global parameters for the match. In this example, there are 40 phenotypes in the gene profile and 22 phenotypes in the evolutionary profile. The overall similarity is 0.23, which is the median of 924 pairwise IC scores. Details of the top pairwise matches are listed below, including the identity of the MICA and the normalized Match IC score. Note that while there were no exact matches between the gene and taxon phenotypes in this example, there are nonetheless a sufficient number of high scoring pairs of phenotypes across upper and lower limbs, such that the overall similarity is higher than would be expected if these profiles had been randomly selected.

**Figure 3.**
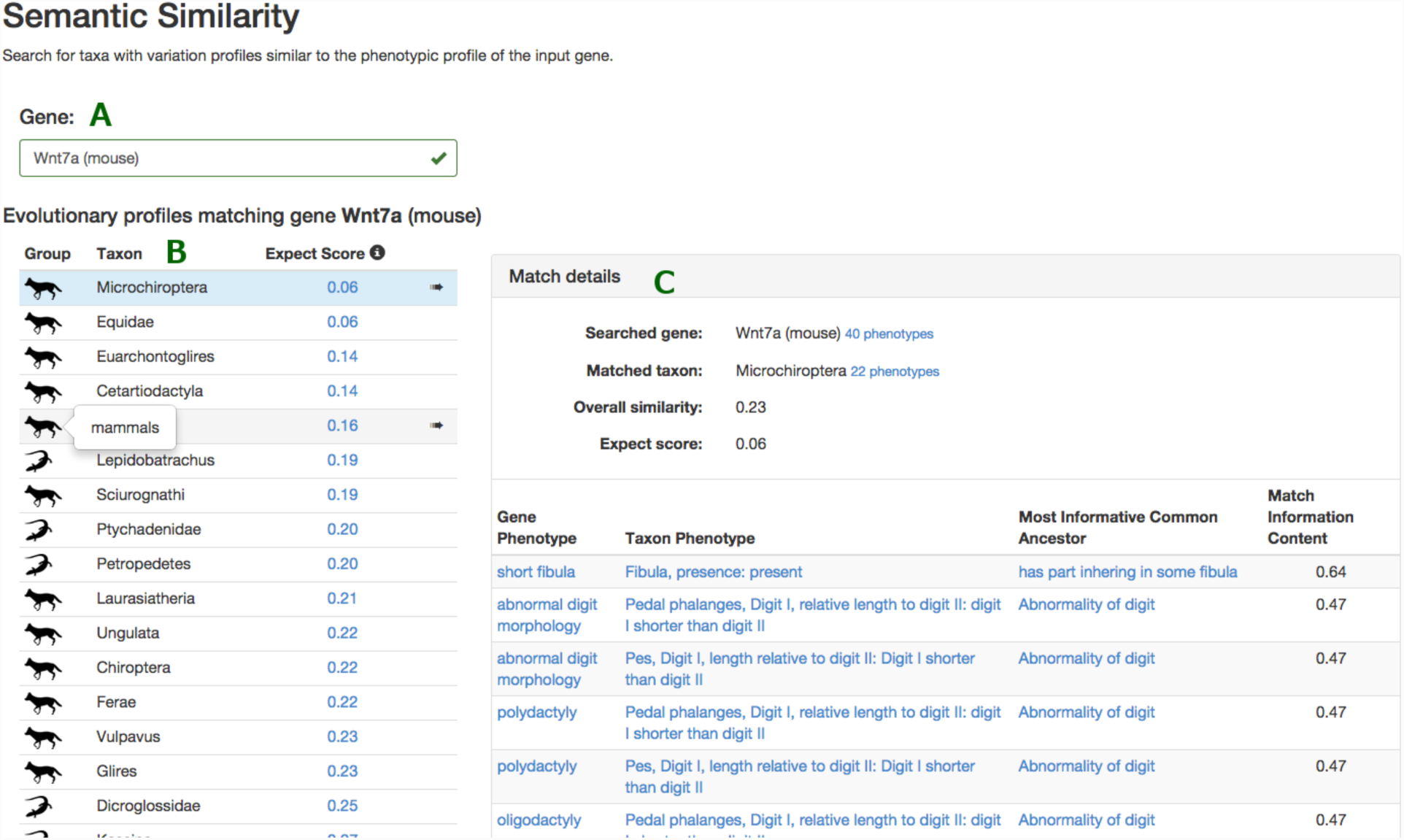
Semantic similarity in the Phenoscape Knowledgebase. (A) Entering a gene name in the search bar brings up a list of genes in model species for which the KB has phenotype annotations. (B) The Summary View shows the 20 evolutionary profiles with the highest Expect Scores (*E*-values) for *Wnt7a* (mouse). The first column shows a pictorial representation (http://phylopic.org/) of a higher group for the matched taxon. For example, the higher group ‘Mammals’ is used to represent the top match for *Wnt7a,* Microchiroptera (Bats). Mouse-over of the graphic displays the higher-level group name. (C) The Match Details View displays the best match. Global parameters are shown at the top; details for the highest-scoring phenotype pairs below.

## Discussion

The Phenoscape KB is one of several recently introduced tools that take advantage of ontologies to search large databases for phenotypically similar entities. Others include Monarch (http://monarchinitiative.org/), PhenomeNET (Hoehndorf et al., 2015), Phenodigm (Smedley et al., 2013), and BOCA (Bauer et al., 2012). Monarch allows users to query using one or more abnormal phenotypes and reports disease models and genes with similar phenotypes. Similarly, PhenomeNET allows users to query diseases, genes, genotypes etc. and reports similar diseases, genes, and experimental drug effects ranked by semantic similarity to the query. Phenodigm conducts semantic similarity comparisons between animal models and human diseases using phenotype information to prioritize reported disease gene candidates. BOCA identifies diseases similar to one or more human phenotypes chosen by a user. These tools collectively demonstrate the power of centralized stores of phenotypic information annotated using common ontologies.

There are a number of subtleties in measuring semantic similarity between collections of phenotypes from different domains that should be taken into account when interpreting results (see also Robinson and Webber 2014). For one, measures based on IC can be sensitive to annotation biases. For instance, the evolutionary phenotype profiles in the Phenoscape KB are more extensive for certain taxonomic groups than others, tend to focus on the skeletal anatomy of fins and limbs, and miss many phenotypes only visible in live specimens. The gene phenotype profiles from the model organism databases have their own sets of biases, including the under-representation of phenotypes with lethal effects and those not observable in whole organisms, and the vagaries of what phenotypes are of interest to individual investigators. The Phenoscape KB calculates IC scores based on the frequencies of evolutionary phenotypes. The search described here will tend to return higher match scores for phenotypes that are rare in the evolutionary dataset, whether that reflects biological rarity or simply annotation bias.

When a strong match cannot be explained away as an annotation bias artifact, it still requires care in biological interpretation. Phenotypic changes resulting from functional evolution in orthologous genes cannot be expected to phenocopy model organism mutations, due to differences in the developmental processes in organisms that are long diverged, the types of genetic changes associated with natural versus induced mutations, the filtering process of natural selection, and the potential for epistatic modifications (see also McGary et al., 2010; Washington et al., 2009). Nonetheless, a strong match is suggestive of some overlap in the affected developmental pathway, and provides a helpful starting point for further empirical investigation.

Other challenges in comparing gene and evolutionary phenotype profiles may be amenable to further methodological refinements. For instance, the differences in coverage between the datasets will lead to many pairwise comparisons for which the MICA in the ontology is unspecific, and thus has a low information content. This limits our ability to identify profiles that have strong similarity for only a subset of phenotypes. Future work is needed explore the suitability of other group-wise semantic similarity measures (Pesquita et al., 2009) to this specific application.

Finally, we wish to emphasize that, in addition to the semantic similarity search described above, the rich content of the Phenoscape KB can be explored in other ways. One can search for information on genes, taxa, characters, or anatomical structures in the KB. One can also perform targeted queries for taxa and filter the results by any phenotype of interest and the individual taxon profiles (without evolutionary inference) can be independently queried.

## Availability

The Phenoscape KB is available at http://kb.phenoscape.org. Source code is available under the MIT license from several repositories under the https://github.com/phenoscape organization, specifically phenoscape-owl-tools for the reasoning pipeline, including semantic similarity computation; phenoscape-kb-services for data services; and phenoscape-kb-ui for the web user interface. The code and a dataset representing a freeze of the semantic similarity search results reported here is available from Zenodo (reasoning pipeline: http://dx.doi.org/10.5281/zenodo.17247; web services: http://dx.doi.org/10.5281/zenodo.17248; web user interface: http://dx.doi.org/10.5281/zenodo.17249).

## Acknowledgments

This work has been supported by grants from the US National Science Foundation (DBI-1062404 and DBI-1062542). We thank C. Mungall for advice on the semantic similarity implementation; W. Dahdul, T.A. Dececchi, N. Ibrahim, and L. Jackson for data curation, and the larger community of ontology contributors and data providers.

